# DGM Is Dead; Long Live DGM-Star! A Quantization-Theoretic Approach to Gamma Rate Discretization

**DOI:** 10.64898/2026.10.30.763052

**Authors:** Roman Sokolovskii, Christoph Stelz, Yuning Wang, Ben Bettisworth, Van Nguyen Hoang, Samuel Martin, Thomas Heinis, Oleksiy M. Kozlov, Bui Quang Minh, Alexandros Stamatakis, Ziheng Yang, Nick Goldman

## Abstract

Modeling substitution rate heterogeneity as a discretized Gamma distribution is standard in phylogenetic inference. Ferretti et al. (2026) have recently demonstrated that the most commonly used equiprobable Gamma rate discretization, known simply as the discrete Gamma model (DGM), substantially overestimates average branch lengths in reconstructed phylogenies when the true rates are drawn from a continuous Gamma distribution. We propose an alternative method to discretize the Gamma rate distribution that considerably reduces the bias in estimated branch lengths and can be used as a drop-in replacement for the standard DGM. The method works especially well on large trees with short branches, yielding up to a 20-fold reduction in bias on simulated phylogenies. It is developed by applying quantization-theoretic tools to the analysis of a simplified star-phylogeny abstraction introduced by Ferretti et al. (2026), where the new method is unbiased by construction. We explain Ferretti et al.‘s counterintuitive observation that increasing the number of rate categories can exacerbate the bias of the standard DGM, even though this should in principle make DGM a better approximation to the continuous case, and offer our method as an immediately available alternative.

## 1 Introduction

Mutation rates vary over genomic sites (Fitch and Margoliash, 1967), with significant consequences for phylogenetic analyses (Yang, 1996). Gamma distribution models describing this rate heterogeneity were first derived from empirical studies (Uzzell and Corbin, 1971) and found effective in early studies (Nei and Gojobori, 1986; Jin and Nei, 1990). In two seminal papers, Yang showed how these models could be incorporated into Felsenstein‘s (1981) maximum likelihood (ML) method for estimation of phylogenetic trees and model parameters: first using the continuous Gamma distribution (Yang, 1993), which remains computationally infeasible for even moderate-size datasets, and then using a discrete approximation of the Gamma distribution (Yang, 1994). This latter method has become a standard in phylogenetic inference, its use alongside a given nucleotide substitution model initially denoted +dG (Yang, 1995; Yang and Roberts, 1995) and later simply as +G or +Γ — the method being so ubiquitous that no indication of the discretization of the Gamma distribution seems generally necessary.

Although there is no mechanistic argument why evolution might favour a Gamma distribution (Uzzell and Corbin, 1971; Felsenstein, 2001), its longevity is likely related to its ability to adopt different shapes (Felsenstein, 2001) even with only one free parameter (the shape, *α*) after the mean rate of substitution is fixed to 1 (Yang, 1996). Yang’s discrete Gamma model (DGM) divides the distribution into *K* rate categories, each with probability 1*/K* and with representative rate given by the mean of the underlying (i.e., continuous) Gamma distribution within that category. This is often denoted by, e.g., +G4 for *K* = 4. Choice of *K* is generally left to the user. There are no degrees of freedom within this discretization process; the rate heterogeneity element of the substitution model depends only on the one parameter *α*, which is generally estimated as part of the ML optimization.

Yang also proposed a nonparametric, finite-mixture, discrete-rates model allowing greater flexibility, whereby the *K* rate categories are assigned arbitrary probabilities (weights) *w*_1_, …, *w*_*K*_ and rates *ρ*_1_, …, *ρ*_*K*_ (Yang, 1995, denotes these by *f*_*i*_ and 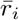). This model, now often denoted FreeRate (Soubrier et al., 2012) or +R, has 2(*K −* 1) free parameters after requiring both the sum of the weights *w*_*i*_ and the mean rate 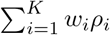 to equal 1. It is common knowledge among developers of phylogenetic inference programs that FreeRate is difficult to implement in a numerically stable manner, due to its increased number of parameters and numerical optimization issues (e.g. Kalyaanamoorthy et al., 2017; Morel et al., 2021).

Yang’s DGM is currently the *de facto* standard for modeling rate heterogeneity over sites. Recently, however, Ferretti et al. (2026) proposed that it should be “retired from use”, that FreeRate might be used instead where possible, and that further work is needed to produce a robust alternative where FreeRate is not suitable. These suggestions were motivated by their observation that branch lengths estimated using DGM can be “biased and sometimes considerably so”. They demonstrated overestimation for *K* = 4 categories and Gamma shape parameter *α* = 0.5, with different phylogenies, and focusing on mean branch lengths or patristic distances between tree tips. Despite this somewhat restricted analysis, e.g. excluding consideration of effects on individual branch lengths, estimates of other parameters, other values of *α*, or model goodness-of-fit, it seems likely that the effect they observed would also occur in other settings. Downstream consequences of this overestimation were not examined, but likely include misinterpretation of divergence levels and, particularly if the overestimation is not uniform across the tree but instead variable on a per-branch level, could affect tree topology inference, phylogenetic dating, and phylogeographic methods. Although Ferretti et al. (2026) studied DGM only in the context of ML inference, we note that it is also the standard in Bayesian phylogenetics (see, e.g., Ronquist et al., 2012; Bouckaert et al., 2019; Baele et al., 2025).

Inspired by Ferretti et al.‘s (2026) presentation of evidence suggesting that the cause of the overes-timation was DGM’s ‘equiprobable’ requirement of equal weight 1*/K* for the *K* rate categories, in this paper we propose an alternative way to discretize the Gamma distribution. This is based on the Lloyd-Max algorithm (Max, 1960; Lloyd, 1982), standard in quantization theory, with adaptations derived by extending Ferretti et al.‘s analysis of bias using a star-phylogeny abstraction.

We call our method DGM-Star (DGM⋆), where the ‘Star’ reflects both its derivation from the star-phylogeny abstraction and its availability as a straightforward ‘drop-in’ replacement for the standard DGM. We show that DGM⋆ dramatically reduces the branch length estimation bias previously reported (Ferretti et al., 2026), especially on large trees with short branch lengths, achieving up to a 20-fold reduction in bias on simulated phylogenies. For long ground-truth branch lengths, we demonstrate that DGM, DGM⋆, and FreeRate all exhibit an underestimation bias, a phenomenon we analyze theoretically on the star phylogeny. We employ high resolution quantization theory to explain why DGM’s bias may *increase* as the number of rate categories *K* grows — a counterintuitive behavior Ferretti et al. observed experimentally in (2026, Fig. S2). This is a critical flaw of the standard DGM.

In our opinion, DGM⋆ has the potential to replace DGM as the new *de facto* standard, expediting DGM’s retirement that was called for by Ferretti et al. (2026) but re-affirming Gamma rate discretization as the dominant technique for modeling rate heterogeneity. While there is space for further improvement, especially when it comes to performance on smaller or more divergent datasets, we expect the quantization-theoretic perspective we introduce here to become an integral part of all future analysis of discretized rate heterogeneity models.

## 2 Methods

### 2.1 Inference Under Rate Heterogeneity

Given a multiple sequence alignment (MSA) **D**, phylogenetic ML inference explores the space of tree topologies, branch lengths, and parameters of the nucleotide substitution and rate heterogeneity models to find a combination that maximizes the likelihood of observing **D**. Like Ferretti et al. (2026), we focus specifically on the inferred branch lengths and their relationship with the chosen rate heterogeneity model.

Standard models of rate heterogeneity assume that each column **D**_*i*_ of an MSA **D** evolves with an independently drawn relative substitution rate *x*_*i*_; each *x*_*i*_ is a realization of a random variable *X* with E[*X*] = 1. Under the rate heterogeneity model of Yang (1993) *X ∼* Gamma(*α*), the continuous Gamma distribution with mean 1 and shape parameter *α*; *α* is typically estimated during ML inference.

Inference under *X ∼* Gamma(*α*) is computationally expensive; to accelerate it, Yang‘s DGM (1994) splits the support [0, *∞*) of Gamma(*α*) into *K* regions at the *K*-tiles of the distribution and assumes that *X* is a discrete random variable that equiprobably takes *K* values — the conditional expectations of the *K* regions. Once *K* is specified by the user, the only remaining parameter of the DGM is again the shape parameter *α* — in contrast to the FreeRate model (Yang, 1995; Soubrier et al., 2012) where the *K* rates and their probabilities are treated as free parameters and are jointly optimized (subject to normalization constraints and resulting in 2(*K −* 1) degrees of freedom).

### 2.2 Rate Discretization as a Quantization Problem

Viewed through the lens of quantization theory (Gray and Neuhoff, 1998), using DGM is performing inference with substitution rate modeled as *q*(*X*), a quantized version of *X ∼* Gamma(*α*). A general quantization rule

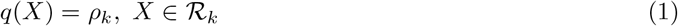

splits the support of *X* into *K* regions, *ℛ*_*k*_ for *k* = 1, … *K*, and assigns to each a corresponding “representative” rate *ρ*_*k*_. The probability *w*_*k*_ of *q*(*X*) = *ρ*_*k*_ is implicit in this formulation and is equal to the probability that *X* falls in the corresponding regio*2n R*_*k*_:

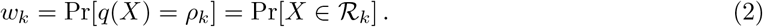

For DGM in particular,

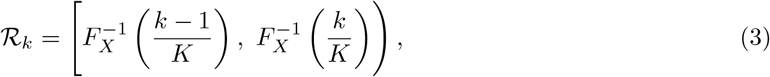

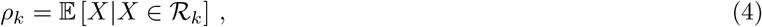

where *F*_*X*_ is the cumulative distribution function of *X ∼* Gamma(*α*), and *w*_*k*_ = 1*/K* by construction.

Ferretti et al. (2026) showed that DGM’s particular choice of *q*(*X*) in (3)–(4) results in biased estimates of average branch lengths. The problem appears to lie in the choice of the regions *R*_*k*_ (3) as opposed to the corresponding representative rates *ρ*_*k*_ (4) since FreeRate also produces (*ρ*_*k*_) that are close to the conditional expectations of the underlying data-generating Gamma distribution without exhibiting DGM’s bias (Ferretti et al., 2026, Fig. 3).

FreeRate abandons attempts to model the discrete rate as *q*(*X*) — i.e., as a quantization of the underlying continuous random variable *X ∼* Gamma(*α*) — altogether and instead models the (*ρ*_*k*_, *w*_*k*_) directly as free parameters. In contrast, we propose an alternative discretization *q*(*X*) to DGM’s equations (3)–(4), which we call DGM⋆. Like standard DGM, our method computes the (*ρ*_*k*_, *w*_*k*_) directly from *α* and can be used as a drop-in replacement for DGM, with *α* also remaining the only parameter to be estimated during ML inference.

### 2.3 DGM⋆

In the framework of quantization theory, *q*(*X*) is designed to optimize a given target function, most commonly minimizing the mean squared error (MSE) E_*X*_ [(*X − q*(*X*))^2^]. In this spirit, DGM can be thought of as designed to maximize *I X*; *q*(*X*), the mutual information between the underlying rate *X* and its quantized version (see Supplementary Sec. 7.1). From the point of view of phylogenetic inference, however, *q*(*X*) should be designed so as to maximize the likelihood of observing **D** given true branch lengths. We believe that it is the mismatch between these two criteria that is the root cause of DGM’s branch length bias, and the deviation of the mean expected substitution rate from 1 under the DGM, demonstrated by Ferretti et al. (2026), is a downstream consequence of that mismatch.

Here, we propose an alternative target function. For a given *α* and assuming *X ∼* Gamma(*α*), DGM⋆ finds a *q*(*X*) that maximizes

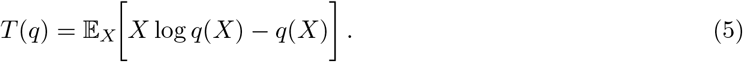

This function emerges from the log-likelihood expression under our extension of the simplified starphylogeny abstraction introduced by Ferretti et al. (2026). As we show in Supplementary Sec. 7.2, maximizing *T* (*q*) yields asymptotically unbiased branch length estimations on the star tree by design.

Although there is no known method to directly calculate the (*ρ*_*k*_, *ℛ*_*k*_) and find *q*(*X*) that maximizes *T* (*q*) (5), we derived an efficient iterative algorithm to do so by adapting the Lloyd-Max algorithm (Max, 1960; Lloyd, 1982) — which targets the MSE and is standard in quantization theory — to target *T* (*q*) instead (Supplementary Sec. 7.3). Supplementary Sec. 7.5 explains how to generate an appropriate starting point of this iteration (asymptotically optimal in the limit of large *K*; see (28)). Inference under DGM⋆ is as fast as under standard DGM (more than 8*×* faster than under FreeRate with the same number *K* = 4 of categories) while achieving lower (i.e., better) BIC scores (Posada and Buckley, 2004) on 78% of the empirical datasets considered (see Sec. 2.4, Supplementary Sec. 7.6, Figs. 5 and 6).

DGM⋆ is available in the latest releases of PAML 4 (Yang, 2007), as option `ncatG=<K> DG-star`; and RAxML-NG 2 (Kozlov et al., 2026) and IQ-TREE 3 (Wong et al., 2026), as option `+Gs` or `+G<K>s`.

### 2.4 Data and Analysis

We use the datasets provided by Ferretti et al. (2026) for our analysis. They ran ML inference on alignments generated from simulated ground-truth trees (we focus on the alignments generated using the continuous Gamma model) and on a subset of 1000 full-genome subtype C HIV-1 sequences available from the PANGEA-HIV consortium (Abeler-Dörner et al., 2019; see Supplementary Sec. 7.6).

In simulations, in addition to using the exact same simulated trees and alignments provided by Ferretti et al. (2026), we generate additional trees using `rtree` in the R `ape` package (Paradis and Schliep, 2019) and alignments using AliSim (Ly-Trong et al., 2023) to explore (*i*) a wider range of tree sizes up to 5000 tips (see Fig. 1C) and (*ii*) average branch lengths up to 10 (see Fig. 1D). Considering larger trees highlights saturation of the bias, and very long branches help explain branch length underestimation in Fig. 2. We use AliSim with options `GTR+FQ+GC{0.5}` to follow Ferretti et al. (2026) and specify a GTR model with equal base frequencies. We note that this combination of `GTR+FQ`, with the exchangeability parameters of GTR not explicitly specified, is in fact equivalent to `JC69` (Jukes and Cantor, 1969). We re-use Ferretti et al.‘s reconstructions under FreeRate where available but re-run reconstructions under DGM for verification; our DGM reconstructions agree with Ferretti et al.‘s (2026). Our analysis of the HIV dataset extends Ferretti et al.‘s procedure to 11 starting 50-sequence ‘seeds’ to alleviate potential seed-dependent bias and combines the results across all seeds.

**Figure 1:**
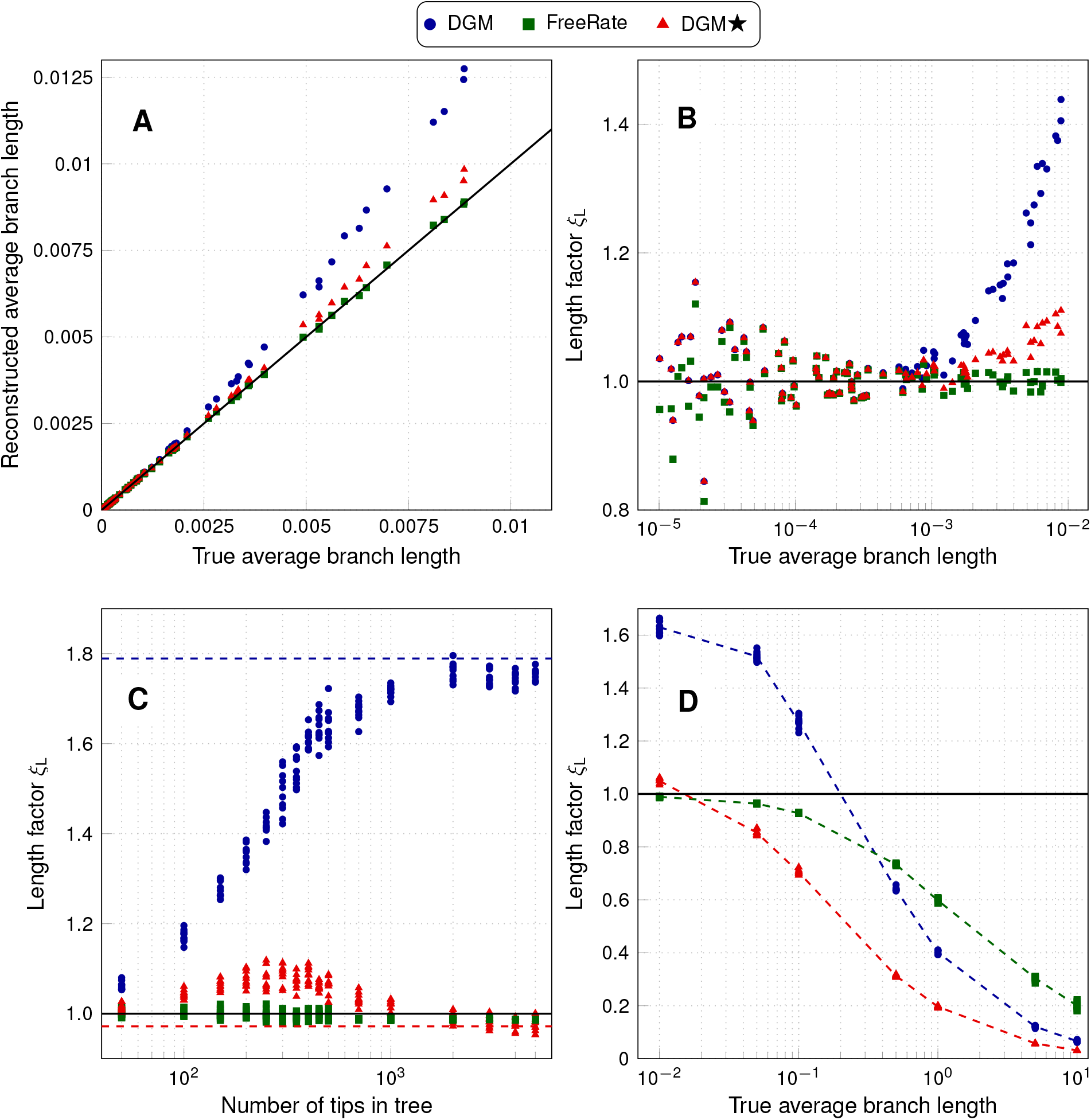
Bias in the average branch length of reconstructed trees with respect to the known ground truth for simulated phylogenies. A continuous Gamma rate distribution with *α* = 0.5 was used to simulate the alignments, each with 25 000 sites. We compare DGM (blue circles), FreeRate (green squares), and DGM⋆ (red triangles) with *K* = 4 discrete rate categories. (A) Reconstructed vs. true average branch length for the 250-tip phylogenies used in Ferretti et al. (2026, Fig. 1A, top right, and Fig. 1B). (B) For the same data as in (A), the ratio of the reconstructed to true average branch length *ξ*_L_ is plotted. (C) *ξ*_L_ as a function of the tree size. The true average branch length is fixed at 0.01. The dashed lines indicate the corresponding *ξ*_L_ on the asymptotic star-phylogeny abstraction in Supplementary Fig. 3 (Supplementary Sec. 7.4) for DGM (blue) and DGM⋆ (red). Each method reconstructs 10 trees of each simulated size. Up to a size of 500 tips, Ferretti et al.‘s data are used (2026, Fig. 1C); we simulated larger trees in an analogous manner. (D) *ξ*_L_ in the long-branch regime. The size of the tree is fixed to 500 tips. Each method reconstructs 10 trees at each simulated true average branch length. The dashed lines connect the empirical means across the 10 replicas.

**Figure 2:**
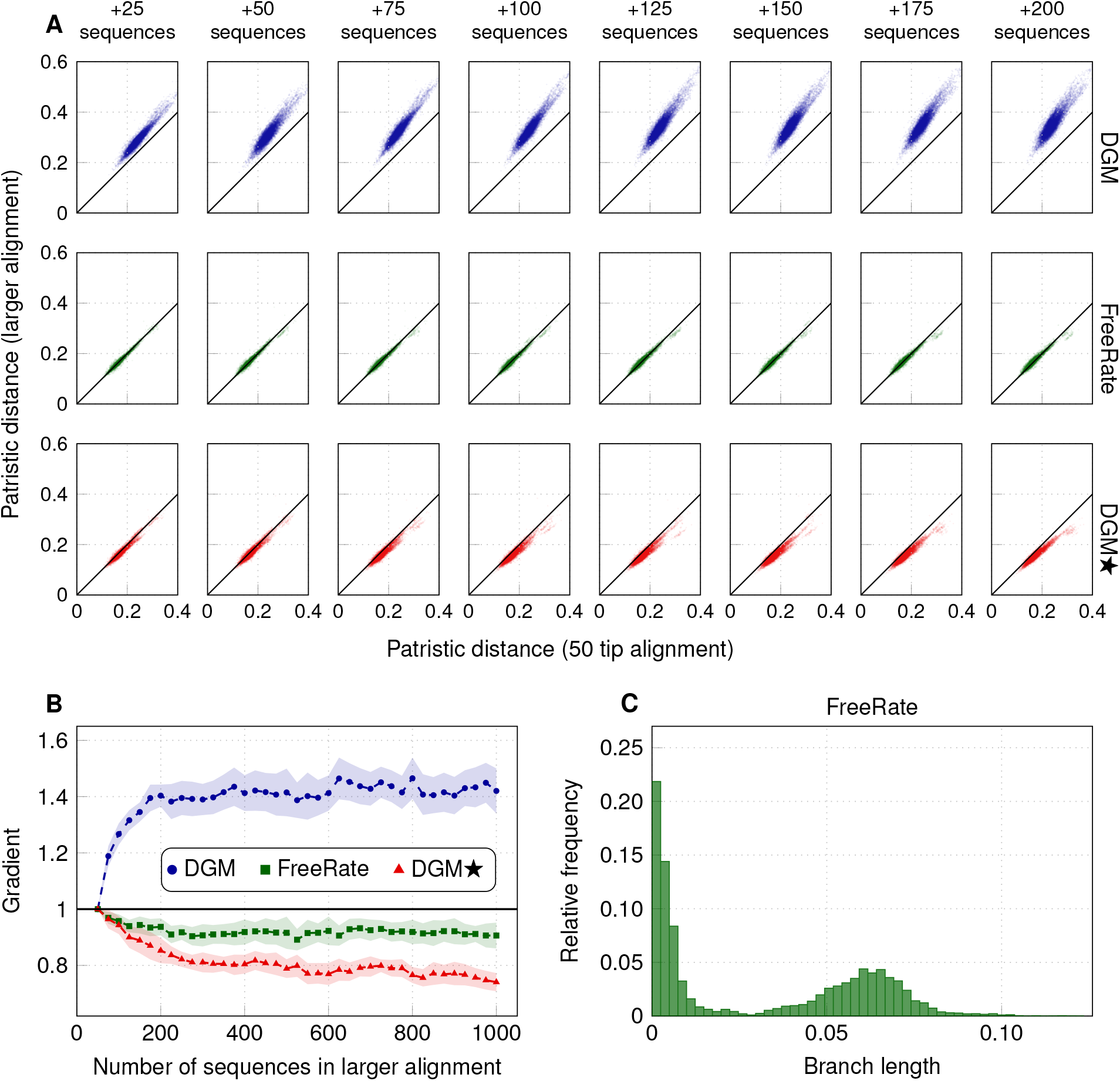
Change in patristic distances as a function of alignment size for the HIV-1 dataset. (A) Pairwise patristic distances between the 50 seeding tips in larger reconstructed phylogenies, plotted against the corresponding patristic distances in the phylogeny reconstructed using only those 50 sequences. Rows, from top to bottom, represent reconstructions using DGM, FreeRate, and DGM⋆, all with *K* = 4 rate categories. Columns represent larger alignments formed by including progressively more sequences in addition to the 50-sequence seed (e.g., the first column uses alignments with 50 + 25 = 75 sequences, where, in addition to the seeding 50 sequences, 25 more sequences were included). The data are merged across 11 samples of 50-sequence seeds, including the seed used by Ferretti et al. (2026) along with their set of progressively larger alignments. (B) The mean slope of the linear regression lines fitted to scatter-point data generated as in (A) but separately for each seed. The shaded areas give the 95% confidence intervals of the gradients across 11 estimations from different seeds. (C) The histogram of branch lengths from the phylogeny inferred by FreeRate using all 1000 HIV-1 sequences.

We use 300 empirical datasets from TreeBASE (Piel et al., 2009) for further performance tests (Supplementary Sec. 7.6). We perform phylogenetic ML inference using RAxML-NG 2 (Kozlov et al., 2026) with *K* = 4 under standard DGM (`GTR+F+G4`), FreeRate (`GTR+F+R4`), and DGM⋆ (`GTR+F+G4s`).

## 3 Results

### 3.1 DGM⋆ Substantially Reduces Bias in Average Reconstructed Branch Lengths

Fig. 1A shows, for a set of simulated phylogenies with 250 tips and alignments generated using the continuous Gamma(*α* = 0.5), the average branch length in the reconstructed tree against the average branch length in the ground-truth tree. Fig. 1B shows the ratio of reconstructed-to-true average branch lengths *ξ*_L_ for the same data (*ξ*_L_ = 1 indicates no bias). The two plots directly mirror Ferretti et al.‘s (2026, Fig. 1A, top right, and Fig. 1B) and were obtained on the same set of simulated trees and alignments. We follow their lead in using *ξ*_L_ to quantify bias, noting that *ξ*_L_ *−* 1 gives the more commonly used signed relative error. DGM⋆ shows a drastically reduced bias compared to the standard DGM; however, the residual bias is higher than FreeRate’s, which does not exhibit any apparent bias in this parameter regime whatsoever.

In Fig. 1C, we investigate how *ξ*_L_ depends on the size of the phylogeny for a fixed ground-truth average branch length of 0.01. This is equivalent to Ferretti et al.‘s (2026) Fig. 1C, extended to explore larger tree sizes. Unlike DGM, which shows *ξ*_L_ growing to reach a plateau of *∼* 1.75, the DGM⋆ bias diminishes as the tree size grows: *ξ*_L_ eventually stabilizes at *∼* 0.97 (red dashed line), corresponding to its asymptotic value on the star tree (see Supplementary Sec. 7.4 and Supplementary Fig. 3).

### 3.2 All Discrete Rate Models Underestimate Very Long Branches

Fig. 1D explores a parameter regime not considered by Ferretti et al. (2026), where the true average branch length is progressively increased, including values beyond the range normally encountered in practical phylogenetic inference. Strikingly, progressively longer branches ultimately lead to a considerable underestimation of average branch lengths for all three methods — including FreeRate, which was not shown to exhibit any substantial bias in Ferretti et al. (2026), and even DGM, which was only shown to overestimate average branch lengths. This behavior aligns with our analysis of the star-phylogeny abstraction in Supplementary Sec. 7.4 (see Supplementary Fig. 3).

### 3.3 HIV-1 Phylogenies Show Signs of Long-Branch Underestimation

We reproduced Ferretti et al.‘s analysis of the HIV-1 dataset (2026, Fig. 2), replicating it across 10 different 50-sequence seeds in addition to the original seed they used (along with their set of progressively larger alignments, i.e., incorporating more sequences); this did not meaningfully affect the results or our conclusions. Fig. 2A compares pairwise patristic distances between the seeding tips in larger phylogenies against the corresponding distances when only the seeding tips are used — the first two rows correspond to DGM and FreeRate and corroborate Ferretti et al.‘s findings (2026, Fig. 2A), while the last row corresponds to our new DGM⋆. Fig. 2B shows the estimated linear regression slopes as a function of the alignment size, which is extended beyond the maximum of 250 tips in Fig. 2A up to the full 1000-sequence dataset (again, the blue curve for DGM corresponds to Ferretti et al.‘s Fig. 2B (2026) while the green curve for FreeRate and the red for DGM⋆ are new). We follow Ferretti et al. (2026) and compare patristic distances in reconstructions from different-sized alignments because here, unlike in Results Sec. 3.1, the ground-truth phylogeny is unknown.

We observe that as the alignment size grows, DGM starts to considerably expand estimated patristic distances by a factor of about 1.4*×* while DGM⋆ contracts them by about 0.8*×*. Surprisingly, FreeRate also shows a systematic shrinkage of patristic distances, by a factor of about 0.9*×*, which was not reported by Ferretti et al. (2026). A representative histogram of branch lengths in Fig. 2C shows that the reconstructed phylogenies contain a substantial fraction of branches of length *∼* 0.05, the value at which rate discretization methods start showing a downward trend in *ξ*_L_ on simulated phylogenies (Fig. 1D) and on the star-phylogeny abstraction (Supplementary Sec. 7.4, Supplementary Fig. 3). Overall, similarly to Results Sec. 3.1 for simulated trees, DGM⋆ shows a smaller bias than standard DGM and a larger bias than FreeRate.

## 4 Discussion

Ferretti et al. (2026) demonstrated that standard DGM shows a substantial average branch length overes-timation bias. (In Supplementary Sec. 7.7, we show that the bias is likely acting at the per-branch level.) As Ferretti et al. observed empirically (2026, HIV-1 results and Figs. S2–S3) and we explain theoretically using the star-phylogeny abstraction (see Supplementary Sec. 7.5 and Supplementary Fig. 4), the bias may first *worsen* as the number of DGM rate categories *K* is increased; we also show that it eventually disappears for *K* asymptotically large. We propose DGM⋆, an alternative Gamma rate discretization procedure that drastically reduces the branch length estimation bias without suffering from FreeRate’s increase in search space complexity.

By design, DGM⋆ works especially well on large trees with short branches (Fig. 1C); for longer branches, the method underestimates branch lengths — a phenomenon also displayed by FreeRate and even DGM (see Results Sec. 3.2, Fig. 1D, and Supplementary Fig. 3). Since long-branch underestimation appears to manifest itself beyond simulations (Results Sec. 3.2) in a real-world HIV-1 dataset (Results Sec. 3.3), alleviating it is a promising direction for future work. One way forward could be to investigate the use of DGM⋆ as a starting point for FreeRate optimization, potentially combining the advantages of both approaches. In addition, our preliminary analysis (not included) shows that DGM⋆ may improve the estimation of the underlying Gamma shape parameter *α*; it would be interesting to explore this further.

There have been other attempts at computationally efficient approximation of the continuous Gamma distribution model of rate heterogeneity. Felsenstein (2001) and Mayrose et al. (2005) investigated Gauss-Laguerre quadrature, which aims to approximate the marginalization of the likelihood over the rate distribution Gamma(*α*) by matching its first 2*K −* 1 moments using the roots of generalized Laguerre polynomials. Felsenstein (2001) shows that as *K* grows, the Gauss-Laguerre quadrature method reaches a plateau in the log-likelihood faster than DGM, indicating that it is a better approximation to the continuous-Gamma integral. However, unlike DGM⋆, the quadrature method is not tailored to any particular “shape” of the likelihood function and is likely to be asymptotically biased for the same reason standard DGM is (see Supplementary Sec. 7.5). An alternative Gauss-Legendre quadrature method (see Yang, 2010) is available in PAML 4 (Yang, 2007). However, since these quadrature methods are rarely used (Pupko and Mayrose, 2020), we leave their detailed analysis for future work.

In summary, while Ferretti et al.‘s call for the retirement of the classic DGM is understandable, our alternative discretization method greatly reduces the bias they described and can be used immediately as a drop-in replacement. We believe that Gamma rate discretization models more generally are likely to remain standard in phylogenetic inference.

It is possible that the quantization-theoretic techniques we employ in this work could prove useful in other areas of phylogenetic inference, beyond modeling mutation rate heterogeneity. Indeed, equiprobable discretization *à la* Yang‘s DGM (1994) has been proposed for acceptance rate ratio (i.e., nonsynonymous/synonymous substitution rate ratio *ω* = *d*_*N*_ */d*_*S*_) by Yang et al. (2000) and recently for diversification rates by Höhna et al. (2026). We make no promises, but we are excited to see future exploration of these and other potential applications of quantization theory in phylogenetic inference under discretized parameter models.

## 5 Data and Code Availability

Alignments, simulated and reconstructed trees used in our analysis, as well as identifiers of PANGEA-HIV sequences and TreeBASE datasets, are available at https://zenodo.org/records/23243178. DGM⋆ is implemented in the latest releases of PAML 4, available at https://github.com/abacus-gene/paml, RAxML-NG 2, available at https://codeberg.org/stelzch/raxml-ng/src/tag/conda/raxml-ng-gamma-2.0.3-h870a6a7_0, and IQ-TREE 3, available at https://github.com/iqtree/iqtree3.

## 6 Acknowledgements

RS is funded via the EU projects DNAMIC 101115389 and NEO 101115317, and received financial support from the European Molecular Biology Laboratory (EMBL) Theory@EMBL Transversal Theme visitor programme. CS is funded by the KiKIT Foundational Research in Helmholtz programme. YW received financial support via Imperial College London’s Undergraduate Research Opportunities Programme. BB and NG are supported by EMBL. VNH is supported by the Australian National University. SM was supported by the Medical Research Council [grant MR/Z503526/1]. TH is supported by Imperial College London. OMK is is funded by the Klaus Tschira Foundation. BQM is supported by the US National Science Foundation Grant DEB-2333243. AS is funded via the EU ERA Chair program under project ID 101087081. ZY is supported by the Biotechnology and Biological Sciences Research Council [grants BB/Y004132/1 and BB/X018571/1].

## 7 Supplementary Material

### 7.1 DGM Maximizes *I* (*X*; *q*(*X*))

To see that the standard equiprobable DGM maximizes the mutual information between the underlying continuous rate *X* and its quantized version *q*(*X*), note that

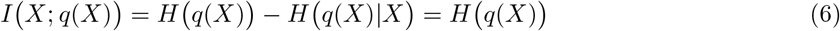

with *H*(*Y*) the Shannon entropy of Y, i.e., − ∑_*y*_ Pr[*Y* = *y*] log Pr[*Y* = *y*]. *H* (*q*(*X*)) is maximized when

*q*(*X*) is uniform.

### 7.2 Deriving the Target Function: the Star Tree Revisited

To derive the target function (5) in Methods Sec. 2.3, we use the simplified star-phylogeny model proposed by Ferretti et al. (2026). The model assumes that *n*_tips_ genomes, each of length *n*_sites_, directly descend from the same root at the center of the resulting “star tree”, and that each of the *n*_tips_ branches of the tree is of length *L*. The genomes comprise binary characters, evolving from the known all-zero genome at the root with an instantaneous site-specific rate *x*_*i*_ for *i* = 1, …, *n*_sites_, each *x*_*i*_ being an independent realization of *X ∼* Gamma(*α*). The probability of observing a substitution 0 *→* 1 over a branch of length *L* and with instantaneous substitution rate *x*_*i*_ is

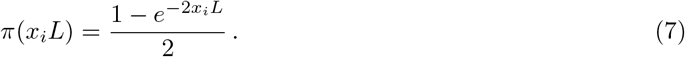

The observed data **D** are the binary genomes at the tips. The inference task is to estimate the branch length *L* by ML ascent — we denote the estimated branch length by 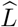.

Crucially, the model used for ML inference does *not* correspond to the above-described data-generating model — specifically, instead of assuming *X ∼* Gamma(*α*), it assumes that *X* takes a set of *K* discrete values *ρ*_*k*_, *k* = 1, …, *K*, each with corresponding probability *w*_*k*_. The goal is to investigate the effect that the choice of (*ρ*_*k*_, *w*_*k*_) has on 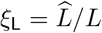.

Ferretti et al. (2026) show that given the number *N*_*i*_ of genomes in **D** that contain a 0 *→* 1 substitution at site *i* = 1, …, *n*_sites_, the log-likelihood *L* of observing **D** under the mismatched rate heterogeneity model is

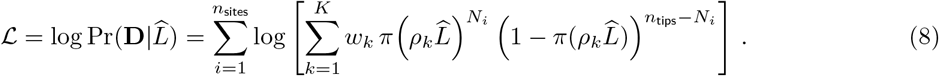

Here, the inner sum is the likelihood of the data observed at each site, marginalized over the discretized rate distribution (*ρ*_*k*_, *w*_*k*_); the outer sum is the log-likelihood of the entire dataset under the assumption that the sites are independent.

We consider the asymptotic limits of long genomes *n*_sites_ *→ ∞*, large trees *n*_tips_ *→ ∞*, and short branches *L →* 0. Of these, the last two were explicitly analyzed by Ferretti et al. (2024); nevertheless, we provide the full derivation here for completeness.

As *n*_tips_ *→ ∞*, the fraction of sites with substitutions *N*_*i*_*/n*_tips_ converges to *π*(*x*_*i*_*L*), the true probability of observing a substitution at site *i* given length *L* and the site-specific realization of the substitution rate *x*_*i*_, and

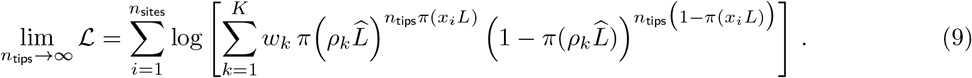

Next, since *x*_*i*_ are independent realizations of *X*, we note that as *n*_sites_ *→ ∞* the outer sum converges to the expectation of the summand with respect to *X* (up to a scaling by *n*_sites_). We also express the summand in terms of the log-sum-exp operator 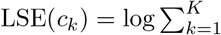, yielding

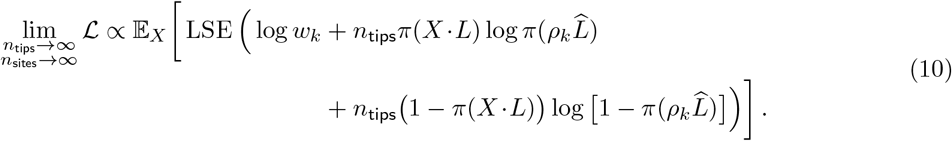

As *n*_tips_ *→ ∞*, LSE(*·*) is dominated by the largest term. After normalization by 1*/n*_tips_, the contribution of the probability term log *w*_*k*_ vanishes, and (10) becomes

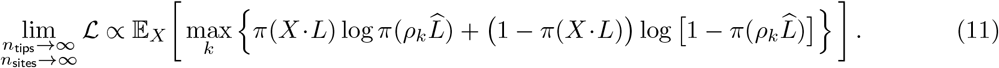

Analogously to (1), we define a quantizer 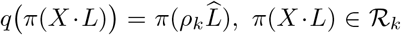, *π*(*X·L*) *∈ ℛ*_*k*_, with the regions *ℛ*_*k*_ chosen as the set of values *π*(*X·L*) for which *k* maximizes the inner expression in (11). Then (11) can be rewritten in terms of *q*(*π*(*X·L*)) as

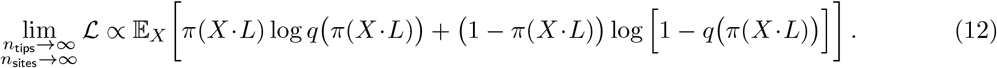

Up to a change of sign, the inner term is the cross-entropy *H*(*p, s*) = *−*E_*p*_[log *s*] of *ℬ* {*q*(*π*(*X · L*))} relative to *ℬ*{*π*(*X · L*))} — two Bernoulli distributions with success probabilities *q* (*π*(*X · L*)) and (*X · L*), respectively — and we can write (12) as

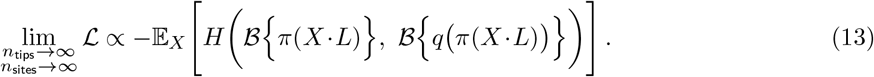

In other words, maximizing the log-likelihood (13) amounts to designing a quantizer *q* (*π*(*X · L*)) that minimizes the expected Bernoulli cross-entropy of *q*( *π*(*X·L*)) relative to *π*(*X·L*).

Optimizing (11)–(13) directly requires access to the distribution of *π*(*X·L*) and therefore cannot be used as a drop-in replacement of the standard DGM. This changes in the limit of small branch lengths, when both *L* and 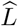 go to zero, where 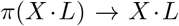 and 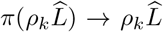. Indeed, substituting these in (11) and ignoring quadratic and higher-order terms in *L* and 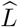, we obtain

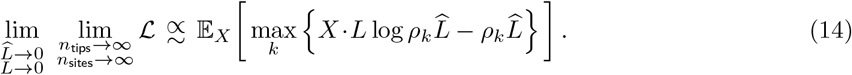

An essentially equivalent expression is derived by Ferretti et al. (2024).

Finally, we apply to (14) the same quantization-theory perspective we used in (12) — instead of fixing (*ρ*_*k*_) and maximizing the log-likelihood *ℒ* as a function of 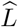, we seek (*ρ*_*k*_) that maximize *ℒ* at 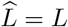 by construction. To do this, we set 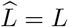 in (14), normalize it by 1*/L* and obtain

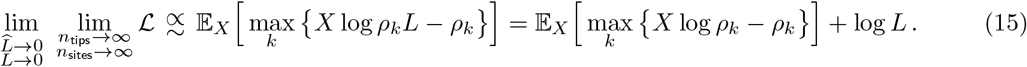

Seeing that the last term does not depend on (*ρ*_*k*_) and using the same quantization-theoretic notation as in (12), we obtain the target function (5):

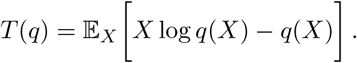

A quantizer *q*(*X*) that maximizes this target function maximizes the likelihood (14) at 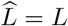.

We emphasize that while Ferretti et al. analyzed the asymptotic limits of *ℒ* in the pre-print version of their paper (Ferretti et al., 2024), they did not recognize the result as the target function for a quantization problem. In essence, while the equiprobable DGM maximizes the mutual information between *X* and *q*(*X*) (Supplementary Sec. 7.1), the correct geometry of the optimization landscape is the Bernoulli cross-entropy (13), which in the limit of short branches becomes the target function *T* (*q*) we propose here (Methods Sec. 2.3).

### 7.3 Deriving Log-Mean Lloyd-Max

For a given target function *T* (*q*) in (5) derived in Supplementary Sec. 7.2, we now seek a procedure to obtain a *q*(*X*) that maximizes it. Specifically, given *K* and *α*, we need to return a set of rates *ρ*_*k*_ and regions *ℛ*_*k*_, *k* = 1, …, *K*, that maximize *T* (*q*) =E_*X*_ [*X* log *q*(*X*) *− q*(*X*)] with *X ∼* Gamma(*α*) and *q*(*X*) defined as in (1).

We achieve this by adapting the Lloyd-Max algorithm (Max, 1960; Lloyd, 1982), a classic result in quantization theory, to target *T* (*q*) instead of the standard MSE. The core idea of the algorithm is to re-calculate the “centroids” *ρ*_*k*_ and their corresponding regions *ℛ*_*k*_ iteratively in a *k*-means-like fashion, alternately fixing one and updating the other until convergence.

With the regions *ℛ*_*k*_ fixed, each *ρ*_*k*_ can be optimized independently of the others; up to terms that do not depend on *ρ*_*k*_, *T* (*q*) becomes

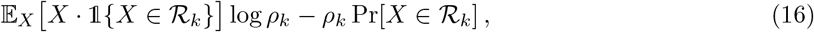

where 1*{·}* is the indicator function. Setting the derivative of (16) with respect to *ρ*_*k*_ to 0, we obtain

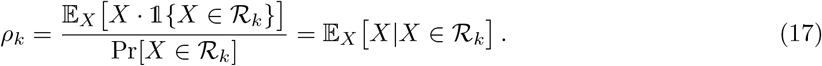

In other words, the centroids *ρ*_*k*_ sit at the conditional expectations of the corresponding regions *ℛ*_*k*_. The same condition must be satisfied in the standard Lloyd-Max when MSE is targeted; furthermore, as Ferretti et al. point out (2026, Fig. 3), standard DGM satisfies it by design, and FreeRate seems to satisfy it empirically.

**Figure 3:**
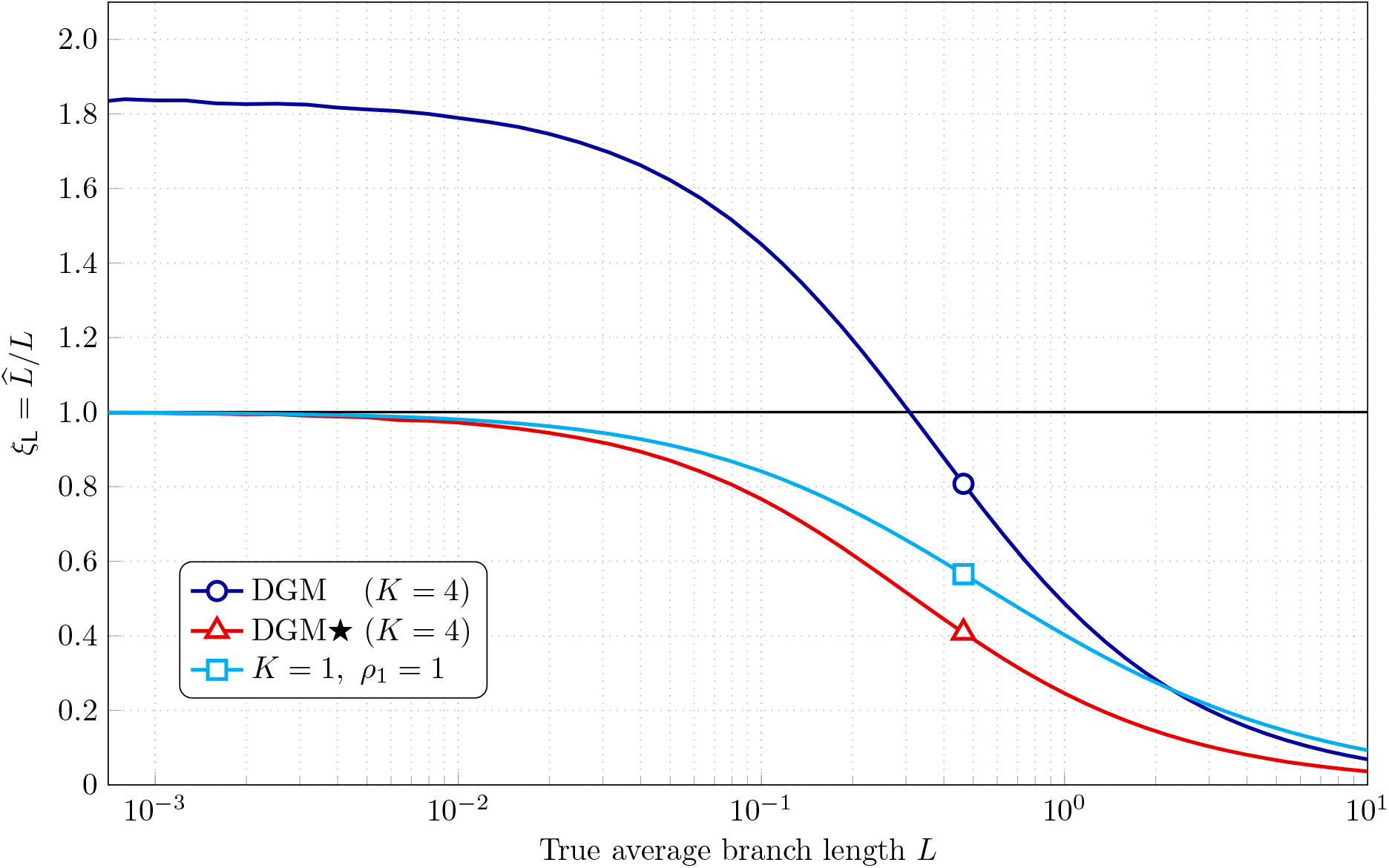
Bias on the asymptotically large star tree (*n*_tips_ *→ ∞, n*_sites_ *→ ∞*) as a function of the true length *L*. The curves are obtained by maximizing *ℒ* in (11) as a function of 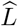.

When it comes to updating the regions *ℛ*_*k*_ given fixed centroids *ρ*_*k*_, our task is to calculate the set of *K −* 1 optimal boundaries *b*_*k*_ between pairs of adjacent centroids (*ρ*_*k*_, *ρ*_*k*+1_) — the *K* regions *ℛ*_*k*_ being [0, *b*_1_), [*b*_1_, *b*_2_), …, [*b*_*K−*1_, *∞*). The key observation is that given a pair of adjacent centroids (*ρ*_*k*_, *ρ*_*k*+1_), the cost of assigning *b*_*k*_ to either region should be the same, yielding

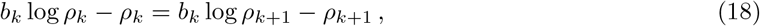

and therefore

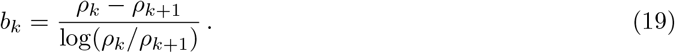

Unlike the standard Lloyd-Max algorithm, where the boundary update rule sets *b*_*k*_ to the arithmetic mean of the two centroids, here the boundary is set to their *logarithmic* mean. Indeed, this is the only change that is needed to adapt the standard Lloyd-Max to target *T* (*q*), and, accordingly, we refer to our modification as the Log-Mean Lloyd-Max algorithm.

Given a starting set of centroids *ρ*_*k*_, Log-Mean Lloyd-Max iterates (19) and (17) until a convergence criterion is met (e.g., the regions *ℛ*_*k*_ change by less than a pre-defined *ϵ*) or a maximum number of iterations is reached. In Supplementary Sec. 7.5, we show how to obtain a good starting set of centroids for the algorithm.

### 7.4 Asymptotic Branch Length Bias on the Star Tree

Fig. 3 plots the asymptotic value of the reconstructed-to-true branch length ratio 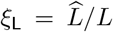 against the true *L*, where 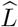 is computed by numerically maximizing (11), for the standard equiprobable DGM (blue, circle marker) and DGM⋆ (red, triangle marker) — both with *K* = 4 — and also for a *K* = 1 discretization with *ρ*_1_ = E_*X*_ [*X*] = 1 (cyan, square marker) for our example of *X ∼* Gamma(*α* = 0.5).

In the limit *L →* 0, DGM exhibits *ξ*_L_ around 1.8 while DGM⋆ is unbiased. Numerical experiments on the star tree, conducted using (8) for a range of finite *n*_tips_ by Ferretti et al. (2026, Fig. 4), show that “no bias in branch lengths is in evidence” for small *L* (Ferretti et al., 2026, Numerical results). Fig. 3 shows, however, that the DGM bias *grows* as *L →* 0. In light of this, Ferretti et al.‘s finite-*n*_tips_ results mean that as *L →* 0, the bias takes progressively larger trees to fully transpire.

**Figure 4:**
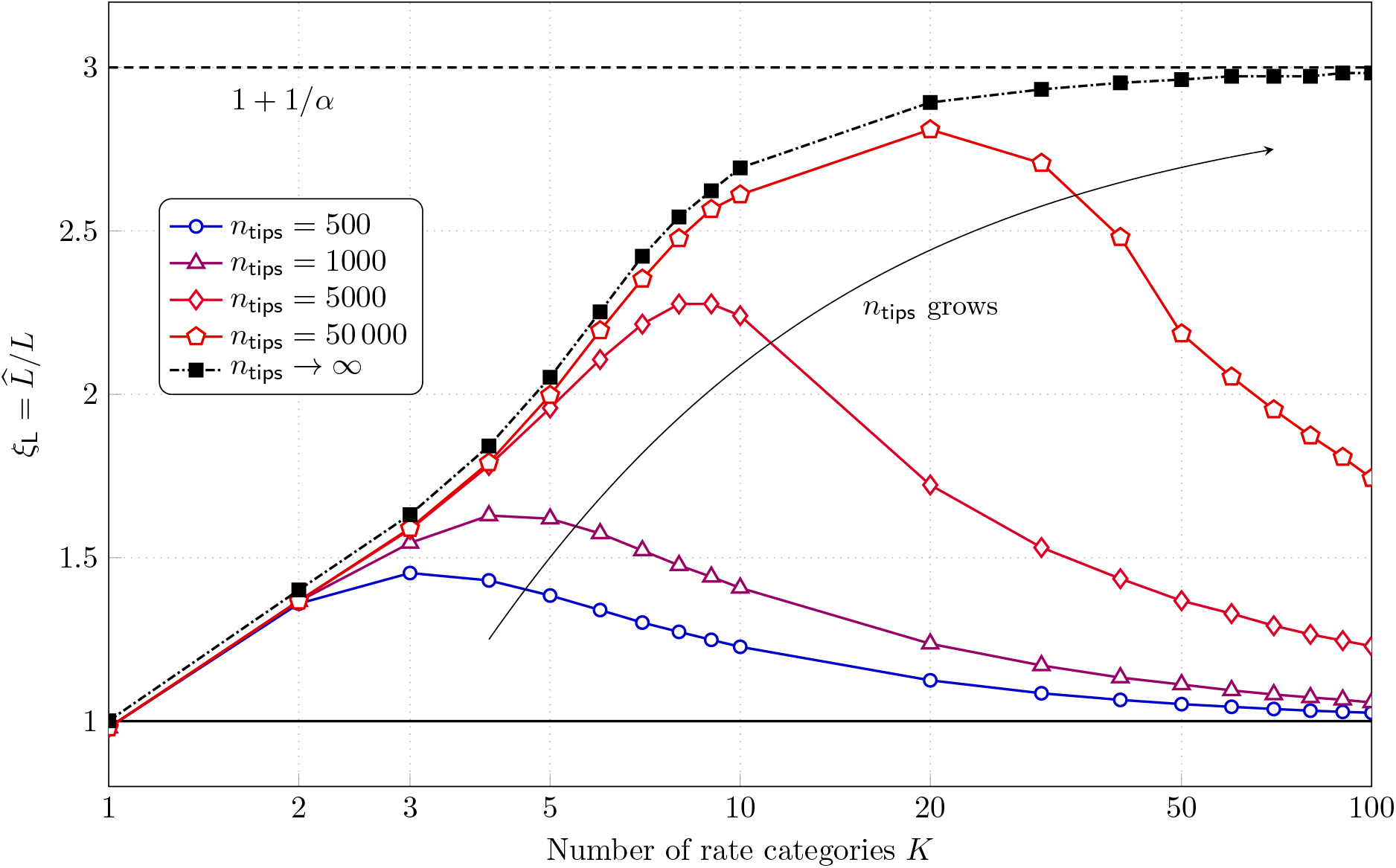
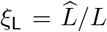 for the standard equiprobable DGM on the star tree with *X ∼* Gamma(*α* = 0.5) as a function of the number of rate categories *K*. Asymptotic case of *n*_tips_ *→ ∞, n*_sites_ *→ ∞*, and *L →* 0 (black, dash-dotted, square markers); finite-length simulations with *n*_sites_ = 30 000, *L* = 0.01, and *n*_tips_ = {500, 1000, 5000, 50 000} (solid, blue to red; circle, triangle, diamond, and pentagon markers, respectively).

Conversely, as *L → ∞*, we observe in Fig. 3 that every method starts severely underestimating *L*. The same phenomenon is present during ML inference on simulated trees (see Fig. 1D). On the star tree and for *K* = 1 (cyan, square marker), it is possible to explain this analytically. In this case the log-likelihood function (13) becomes

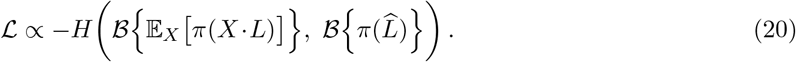

The cross-entropy is minimized when the two distributions are equal, i.e., 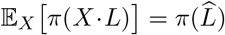, yielding

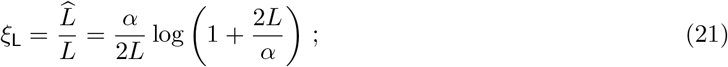

this, in turn, goes to 0 as *L → ∞*. To derive (21), it is useful to recall the moment-generating function of Gamma(*α*), E_*X*_ [*e*^*tX*^ ]= (1 *− t/α*)^*−α*^ for *t < α*.

That one-point discretization *ρ*_1_ = E_*X*_ [*X*] = 1 should underestimate branch lengths for large *L* can be seen from E_*X*_ [*π*(*X· L*)] *≤ π*(E_*X*_ [*X· L*]), a direct consequence of Jensen’s inequality for concave *π*(*·*) — when a certain average fraction of 0 *→* 1 sites is observed, the discretized model will match this at a smaller 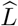 than the true *L*. The same phenomenon is likely in play for *K >* 1, and developing a method to correct the branch length bias for longer *L* is an interesting direction for future work, especially since empirical data show some evidence of such underestimation (see Results Sec. 3.3 and Fig. 2B). (It should be noted, however, that in the limit *L → ∞* information about *L* and *α* is erased from **D**, and their robust estimation becomes fundamentally impossible.)

### 7.5 Increasing the Number of Categories May *Hurt* DGM: an Analysis on the Star Tree

Ferretti et al. observed empirically (2026, in their HIV-1 results and Figs. S2 and S3) that increasing *K* may lead to more-inflated patristic distances (see Results Sec. 3.3), probably caused by more-biased estimations of branch lengths. This seems surprising, as we may imagine that increasing *K* would make DGM a better approximation to the continuous case, reducing the model misspecification that lies at the heart of the average branch length estimation bias. Here, we explain this phenomenon on the star tree. First, we leverage high-resolution quantization theory to show that DGM is biased in the limit of small *L* and large *n*_tips_, *n*_sites_, and *K*. Second, we show that when *n*_tips_ is fixed and finite, DGM is *unbiased* for large *n*_sites_ and *K*. Last, we demonstrate numerically how the tension between the two incompatible asymptotic regimes is reconciled at finite parameter values.

#### 7.5.1 Asymptotically Large *n*_tips_ and *n*_sites_: the Bias Worsens as *K* Grows

In Fig. 4, the black dash-dotted curve with square markers shows 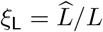 as a function of *K*, estimated by maximizing (14) for standard equiprobable DGM and *X ∼* Gamma(*α* = 0.5). (Note that (14) assumes *n*_tips_ *→ ∞, n*_sites_ *→ ∞*, and *L →* 0.) We observe that *ξ*_L_ first grows sharply with *K* and then plateaus at *ξ*_L_ *≈* 3. In the following, we prove that in fact in this regime *ξ*_L_ tends to exactly 3 as *K → ∞* for *α* = 0.5, and more generally *ξ*_L_ *→* 1 + 1*/α* for *α <* 1.

First, we introduce a distortion function

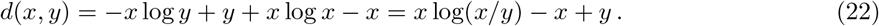

Note that, since E_*X*_ [*X* log *X − X*] does not depend on *q*(*X*), minimizing E_*X*_ [*d*(*X, q*(*X*)] is equivalent to maximizing the target function *T* (*q*) from (5). The distortion *d*(*x, y*) is locally quadratic, i.e., *d*(*x, x*) = 0 and

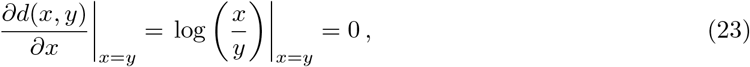

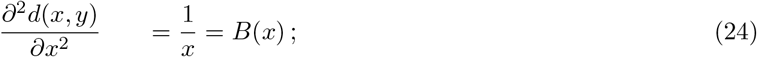

these three conditions mean that for *x ≈ y, d*(*x, y*) is well approximated as

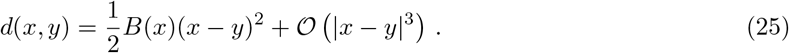

Let *λ*_*q*_(*x*) be the asymptotic proportion of quantization levels around *x* for a quantizer *q* in the limit *K → ∞*, referred to as the *point density* of *q*. It is a known result in high-resolution quantization theory (Gardner and Rao, 1995; Gray and Neuhoff, 1998) that for locally quadratic *d*(*x, y*) the optimal point density *λ*^*\**^(*x*) satisfies

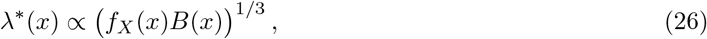

where *f*_*X*_ (*x*) is the probability density function (PDF) of X. In our case of (22)–(24) and *X ∼* Gamma(*α*), this amounts to

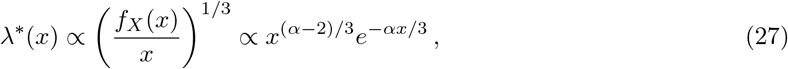

and therefore *λ*^*\**^(*x*) is the PDF of the Gamma distribution with shape parameter *α*^*′*^ = (*α* + 1)*/*3 and rate parameter *β* = *α/*3.

As a useful aside, we note that the knowledge of the optimal point density *λ*^*\**^(*x*) allows us to obtain a good starting point for the Log-Mean Lloyd-Max algorithm by setting

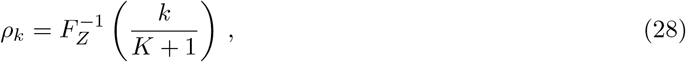

where *F*_*Z*_(*·*) is the cumulative distribution function of *Z ∼* Gamma(*α*^*′*^, *β*). This technique is known as *companding* in quantization theory (Gray and Neuhoff, 1998). The density of *ρ*_*k*_ generated as in (28) tends to *λ*^*\**^(*x*) as *K → ∞* and therefore is asymptotically optimal.

Now, consider the point density *λ*_E_(*x*) of the equiprobable DGM quantizer *q*_E_(*X*). For each quantization region to have the same probability, as must be the case for *q*_E_(*X*), the density of regions should compensate for probability density, so *λ*_E_(*x*) = *f*_*X*_ (*x*). Evaluating the log-likelihood expression (14) at 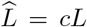 amounts to multiplying each *ρ*_*k*_ in (15) by *c* — or, equivalently, to using a scaled quantizer *q*_*c*_(*X*) = *c · q*_E_(*X*) in (5). (The value of *c* that maximizes (5) for *q*(*X*) = *q*_*c*_(*X*) determines the observed bias *ξ*_L_.) The point density of *q*_*c*_(*X*) is

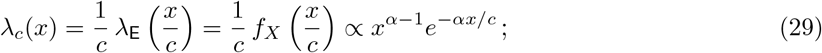

comparing (27) with (29) for *α* = 0.5 reveals that *λ*_*c*_(*x*) = *λ*^*\**^(*x*) for *c* = 3, so asymptotically *ξ*_L_ *→ c* = 3.

To derive the likelihood-maximizing *c* more generally as a function of *α*, we first write the Bennett’s integral (Bennett, 1948) for locally-quadratic distortions (Gardner and Rao, 1995; Gray and Neuhoff, 1998),

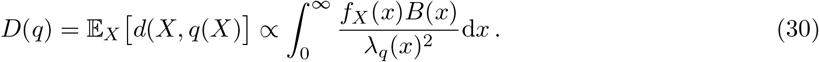

For our *X ∼* Gamma(*α*), *λ*_*q*_(*x*) = *λ*_*c*_(*x*) from (29), and *B*(*x*) = 1*/x* from (24),

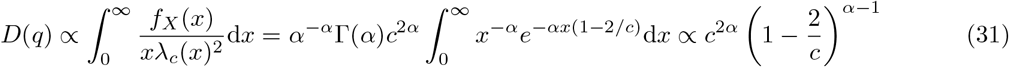

up to terms that do not depend on *c* and for *α <* 1. Setting *∂D*(*q*)*/∂c* = 0, we obtain from (31) that

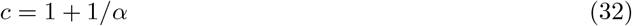

for *α <* 1. For *α ≥* 1, the distortion integral (31) diverges.

In summary, on the asymptotically large star tree with *n*_tips_ *→ ∞, n*_sites_ *→ ∞*, and small branch lengths *L →* 0, *ξ*_L_ grows with *K* towards the asymptotic limit of 1 + 1*/α*. Imagine that in practice a high-stakes analysis of a large alignment with long genomes is performed. Hoping for robust inference, standard DGM with a large *K* is chosen in the expectation that this guarantees a good fit of the discretized Gamma to the underlying continuous version, despite incurring the corresponding cost in increased computational complexity. Our analysis in this section shows that in fact the decision to choose a higher *K* may lead to a *more* biased estimation of branch lengths — a critical problem for standard equiprobable DGM.

#### 7.5.2 Fixed *n*_tips_ and Growing *n*_sites_: the Bias Disappears as *K* Grows

We now consider a different asymptotic regime where *n*_sites_ *→ ∞* and *K → ∞* as before, but *n*_tips_ is fixed and finite. This is an important regime because here the frequency of each of the 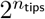 possible alignment columns converges to its true probability, and unbiased inference is a good indication of statistical consistency of a model. In the following, we show that in this setup standard DGM is asymptotically unbiased.

Recall that in the simplified star-phylogeny model, the log-likelihood can be expressed as (8). For the standard equiprobable DGM,

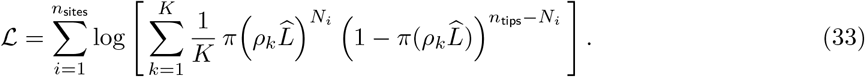

In the limit *K → ∞*, this becomes

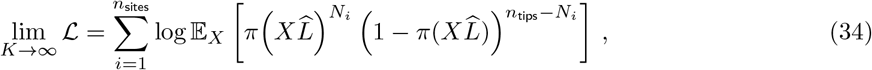

amounting to inference under the true data-generating model of rate heterogeneity. As *n*_sites_ *→ ∞*, the outer sum also tends to an expectation (up to 1*/n*_sites_),

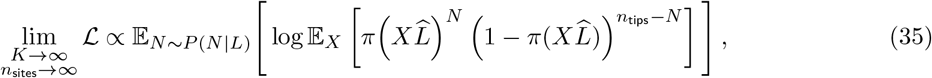

where the distribution of the number of substitutions per site *N* is given by a marginalization

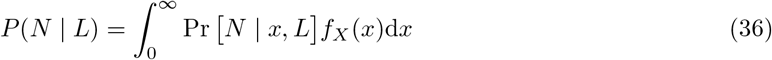

with the true underlying relative substitution rate *X ∼* Gamma(*α*) and *N* | *X, L ∼* Binomial *n*_tips_, (*π*(*X·L*)) .

Similarly, the inner expectation over *X* can also be expressed as a marginalization,

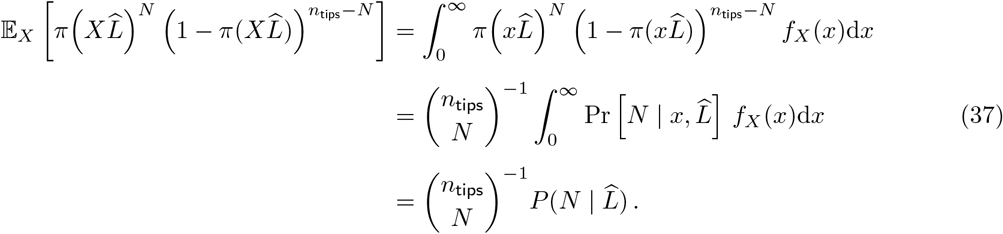

Substituting this result into (35) yields

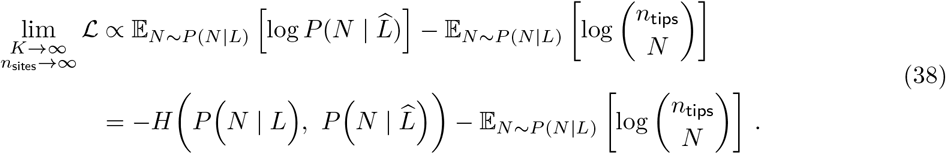

The second term does not depend on 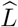, and the negative cross-entropy term (and therefore the log-likelihood) is maximized when 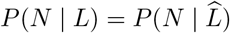 — i.e., when 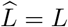.

#### 7.5.3 Finite *n*_tips_, *n*_sites_, and *K*: Squaring the Circle

In Supplementary Sec. 7.5.1, we show that on the asymptotically large star tree *ξ*_L_ tends to 1 + 1*/α* as *K → ∞*. In contrast, Supplementary Sec. 7.5.2 proves that as *K → ∞* the bias *disappears* (i.e., *ξ*_L_ *→* 1) for any fixed *n*_tips_. What does it mean for finite *n*_tips_, *n*_sites_, and *K*?

Fig. 4 shows, alongside the asymptotic large-star *ξ*_L_ curve (black, dash-dotted, square markers), simulated *ξ*_L_ curves for *L* = 0.01, *n*_sites_ = 30 000, and *n*_tips_ = *{*500, 1000, 5000, 50 000*}* against *K* (solid, blue to red; circle, triangle, diamond, and pentagon markers, respectively). Each dot of the finite-*n*_tips_ curves is obtained from 500 Monte Carlo-simulated realizations of *N*_*i*_ by maximizing (8) to estimate 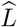 and taking the mean. We observe that the simulated curves exhibit a pull towards both asymptotes: for large *K* they are eventually brought down towards unbiased estimation (*ξ*_L_ = 1, black solid line) in agreement with Supplementary Sec. 7.5.2, but first they rise alongside the large-star curve (black dash-dotted line) and towards an increasing *ξ*_L_ *→* 1 + 1*/α* (black dashed line) as analyzed in Supplementary Sec. 7.5.1. For progressively larger *n*_tips_ the pull towards the black dash-dotted curve gets stronger, and it takes a larger *K* to bring it down. This is bad news for the standard DGM: it means that progressively larger *K* are required for unbiased estimation precisely when increasing *K* becomes less affordable.

### 7.6 Evaluating DGM⋆ on Empirical Datasets

We used the same alignment of 1000 full-genome subtype C HIV-1 sequences as Ferretti et al. (2026) and applied their filtering procedure, removing positions where more than 10% of sequences had an `N` or gap character. We obtained an alignment with 7838 sites. While this does not match the sequence length of 7854 sites reported by Ferretti et al. (2026), the difference appears negligible and no consequences were detectable in our downstream analyses.

Additionally, we evaluated DGM, DGM⋆, and FreeRate with *K* = 4 categories on 300 empirical datasets from TreeBASE (Piel et al., 2009) (median 212 taxa, mean 27 835 alignment sites). The runtime of evaluating all datasets under DGM⋆ was on par with DGM (slowdown of 1.01*×*), whereas FreeRate took 8.35*×* as long (Fig. 5). On 234 datasets (78%), DGM⋆ achieved a lower BIC score than DGM (Fig. 6), indicating better model fit (Posada and Buckley, 2004).

**Figure 5:**
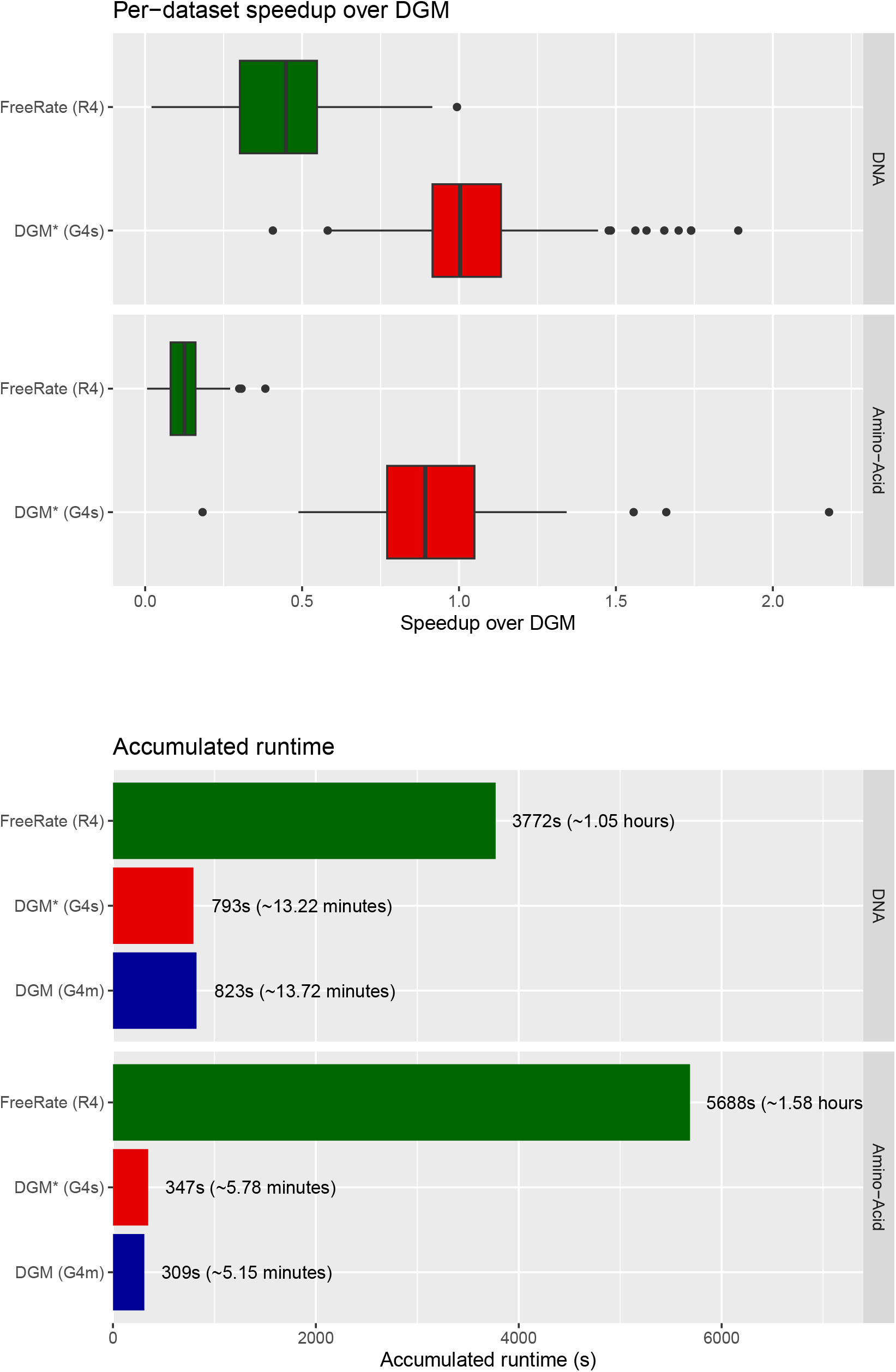
Runtime performance comparison on TreeBASE. Top: per-dataset speedups relative to DGM. Bottom: total runtime over all datasets, with DNA and amino acid datasets reported separately.

**Figure 6:**
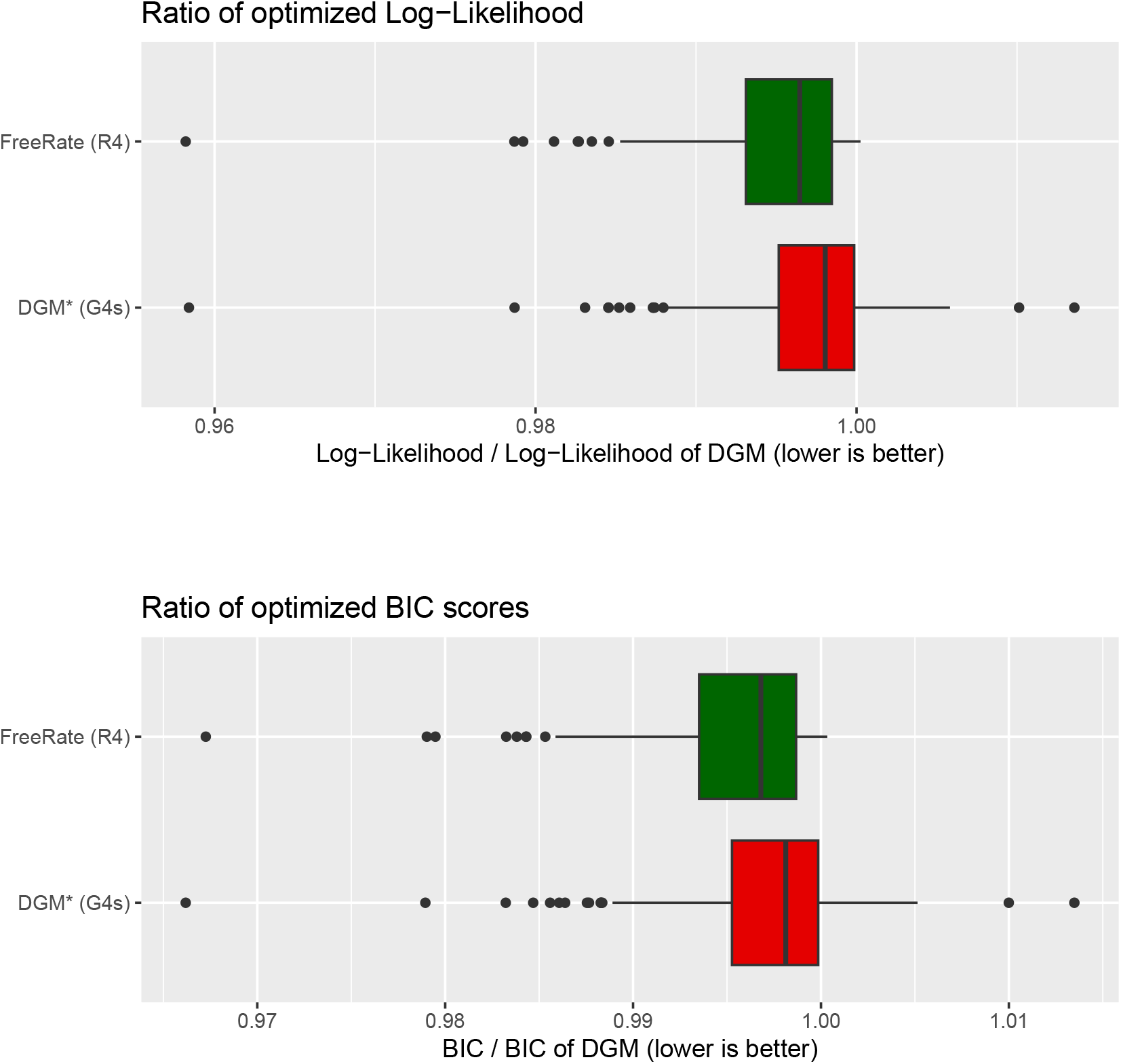
Log-likelihood (top) and BIC (bottom) comparison of FreeRate and DGM⋆ relative to standard DGM on the 300 TreeBASE datasets.

Benchmarks were performed on an AMD Ryzen 9 3950X 16-core processor with 8 threads, using the conda package `stelzch::raxml-ng-gamma-2.0.3-h870a6a7 0`, AlmLinaux 10.2, Kernel 6.12.0-211.51.1.el10 2.x86 64. Speedups are computed using the mean of 3 repetitions per dataset and the RHAS model. Results in this section were corroborated using IQ-TREE 3 (Wong et al., 2026) (details omitted for brevity).

### 7.7 Branch Length Bias Acts on Each Branch Individually

In Supplementary Sec. 7.4, Fig. 3 showed how 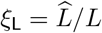 varies with true branch length *L* on the star tree, where there is an asymptotically decreasing trend towards 0. The corresponding trend in the average branch lengths of simulated complex phylogenetic trees is shown in Fig. 1D. Here, we investigate whether the bias affects the entire tree as a whole or rather each branch individually in a length-dependent manner.

Fig. 7 shows the ratio *ξ*_L_ of the individual tip (i.e., terminal) branch lengths inferred by DGM over the true tip branch lengths for a single simulated 500-tip tree. The simulated tree is designed by generating branches with lengths evenly distributed in log-space across a wide range between 10^*−*4^ and 10^2^. (Note that we omit points that correspond to the 23 branches of reconstructed length 10^*−*6^ — the default minimum reconstructed branch length in RAxML-NG 2.)

**Figure 7:**
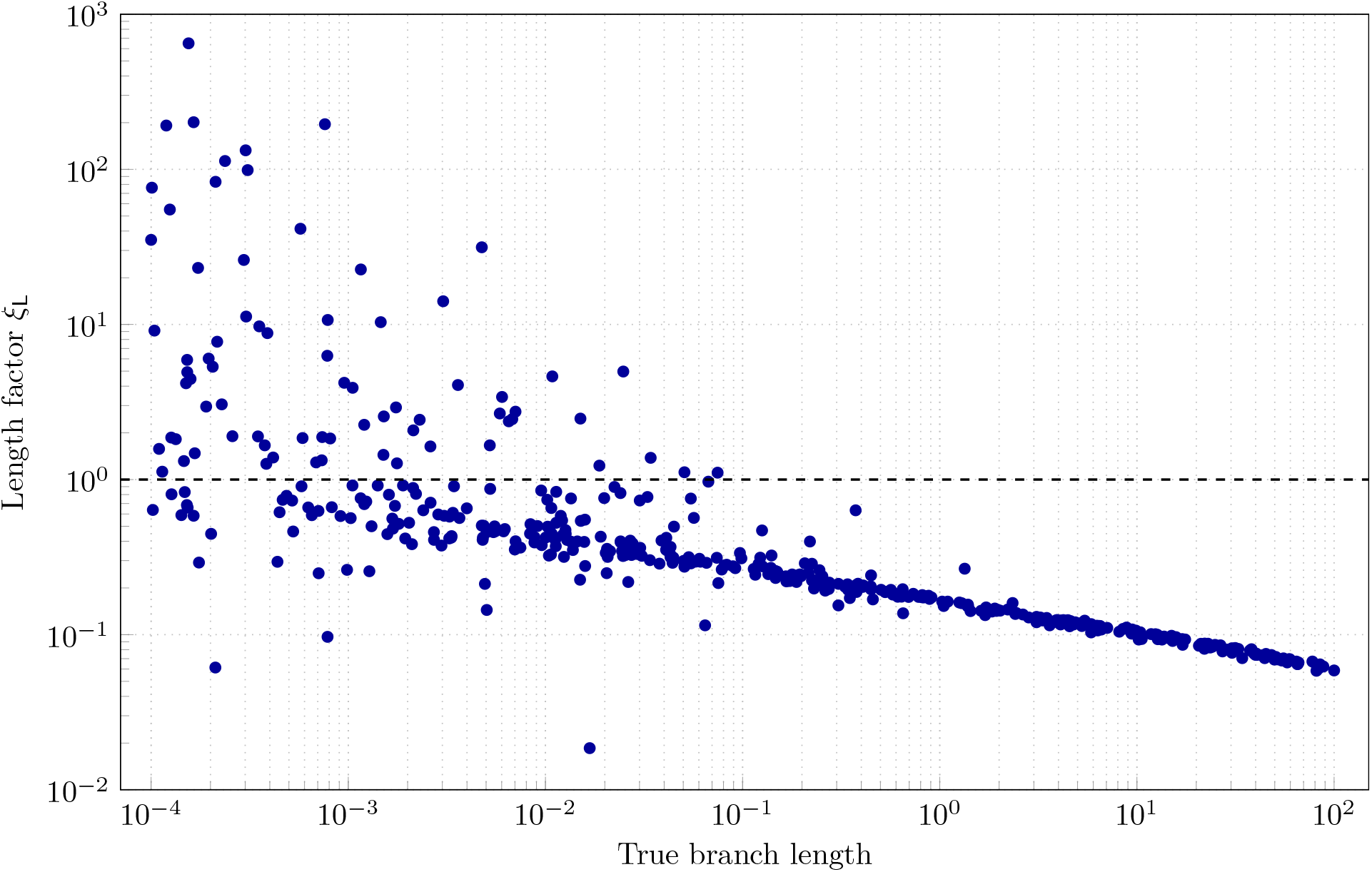
*ξ*_L_ under standard DGM for individual tip branches as a function of the true tip branch lengths on a simulated 500-tip tree with branch lengths distributed evenly in log-space between 10^*−*4^ and 10^2^. The points are obtained by matching corresponding tips from the inferred tree and the true tree.

For short branch lengths, the values of *ξ*_L_ are widely distributed with some fraction of tip branch lengths being overestimated (*ξ*_L_ *>* 1). As the true branch lengths grow, the variation in *ξ*_L_ decreases, and the value of *ξ*_L_ itself decreases with it. In other words, the underestimation increases with increasing branch length, approximately proportionately to the true branch length raised to the power 1/4 (the approximate value of the downward gradient of the points in log-log space). Crucially, this means that *ξ*_L_ varies within a single tree, suggesting that the bias acts on the per-branch level.

Running FreeRate and DGM⋆ on the same simulated tree shows a similar downward trend in *ξ*_L_ towards progressively more underestimation as the branch length grows (not shown), aligning with the analysis in Supplementary Sec. 7.4 where every discretization method exhibits severe underestimation in the limit of *L → ∞*.

## References

Abeler-Dörner, L., Grabowski, M. K., Rambaut, A., Pillay, D., and Fraser, C. (2019). PANGEA-HIV 2: phylogenetics and networks for generalised epidemics in Africa. Current Opinion in HIV and AIDS, 14:173–180.

Baele, G., Ji, X., Hassler, G. W., McCrone, J. T., Shao, Y., Zhang, Z., Holbrook, A. J., Lemey, P., Drummond, A. J., Rambaut, A., and Suchard, M. A. (2025). BEAST X for Bayesian phylogenetic, phylogeographic and phylodynamic inference. Nature Methods, 22:1653–1656.

Bennett, W. R. (1948). Spectra of quantized signals. The Bell System Technical Journal, 27:446–472.

Bouckaert, R., Vaughan, T. G., Barido-Sottani, J., Duchêne, S., Fourment, M., Gavryushkina, A., Heled, J., Jones, G., Kühnert, D., De Maio, N., Matschiner, M., Mendes, F. K., Müller, N. F., Ogilvie, H. A., du Plessis, L., Popinga, A., Rambaut, A., Rasmussen, D., Siveroni, I., Suchard, M. A., Wu, C.-H., Xie, D., Zhang, C., Stadler, T., and Drummond, A. J. (2019). BEAST 2.5: an advanced software platform for Bayesian evolutionary analysis. PLOS Computational Biology, 15:e1006650.

Felsenstein, J. (1981). Evolutionary trees from DNA sequences: a maximum likelihood approach. Journal of Molecular Evolution, 17:368–376.

Felsenstein, J. (2001). Taking variation of evolutionary rates between sites into account in inferring phylogenies. Journal of Molecular Evolution, 53:447–455.

Ferretti, L., Golubchik, T., Di Lauro, F., Ghafari, M., Villabona-Arenas, J., Atkins, K. E., Fraser, C., and Hall, M. (2024). Biased estimates of phylogenetic branch lengths resulting from the discretised Gamma model of site rate heterogeneity. bioRxiv. doi:10.1101/2024.08.01.606208.

Ferretti, L., Golubchik, T., Di Lauro, F., Ghafari, M., Villabona-Arenas, J., Atkins, K. E., Fraser, C., and Hall, M. (2026). Biased estimates of phylogenetic branch lengths resulting from the discretised Gamma model of site rate heterogeneity. Systematic Biology, syag037. doi:10.1093/sysbio/syag037.

Fitch, W. M. and Margoliash, E. (1967). A method for estimating the number of invariant amino acid coding positions in a gene using cytochrome c as a model case. Biochemical Genetics, 1:65–71.

Gardner, W. and Rao, B. (1995). Theoretical analysis of the high-rate vector quantization of LPC parameters. IEEE Transactions on Speech and Audio Processing, 3:367–381.

Gray, R. M. and Neuhoff, D. L. (1998). Quantization. IEEE Transactions on Information Theory, 44:2325–2383.

Höhna, S., Freyman, W. A., Nolen, Z., Huelsenbeck, J. P., May, M. R., Rannala, B., and Moore, B. R. (2026). Inferring branch-specific rates of lineage diversification under the birth–death-shift process. Systematic Biology, 75:948–966.

Jin, L. and Nei, M. (1990). Limitations of the evolutionary parsimony method of phylogenetic analysis. Molecular Biology and Evolution, 7:82–102.

Jukes, T. H. and Cantor, C. R. (1969). Evolution of protein molecules. In Munro, H. N., editor, Mammalian Protein Metabolism, chapter 24, pages 21–132. Academic Press.

Kalyaanamoorthy, S., Minh, B. Q., Wong, T. K. F., von Haeseler, A., and Jermiin, L. S. (2017). Modelfinder: fast model selection for accurate phylogenetic estimates. Nature Methods, 14:587–589.

Kozlov, O. M., Togkousidis, A., Stelz, C., Höhler, D., Wiegert, J., and Stamatakis, A. (2026). RAxML-NG 2: automatic model selection, novel tree search heuristics, and fast branch support metrics. bioRxiv. doi:10.64898/2026.09.09.750097.

Lloyd, S. (1982). Least squares quantization in PCM. IEEE Transactions on Information Theory, 28:129–137.

Ly-Trong, N., Barca, G. M. J., and Minh, B. Q. (2023). AliSim-HPC: parallel sequence simulator for phylogenetics. Bioinformatics, 39:btad540.

Max, J. (1960). Quantizing for minimum distortion. IRE Transactions on Information Theory, 6:7–12.

Mayrose, I., Friedman, N., and Pupko, T. (2005). A Gamma mixture model better accounts for among site rate heterogeneity. Bioinformatics, 21:ii151–ii158.

Morel, B., Barbera, P., Czech, L., Bettisworth, B., Hübner, L., Lutteropp, S., Serdari, D., Kostaki, E.-G., Mamais, I., Kozlov, A. M., Pavlidis, P., Paraskevis, D., and Stamatakis, A. (2021). Phylogenetic analysis of SARS-CoV-2 data is difficult. Molecular Biology and Evolution, 38:1777–1791.

Nei, M. and Gojobori, T. (1986). Simple methods for estimating the numbers of synonymous and nonsynonymous nucleotide substitutions. Molecular Biology and Evolution, 3:418–426.

Paradis, E. and Schliep, K. (2019). ape 5.0: an environment for modern phylogenetics and evolutionary analyses in R. Bioinformatics, 35:526–528.

Piel, W. H., Chan, L., Dominus, M. J., Ruan, J., Vos, R. A., and Tannen, V. (2009). TreeBASE v. 2: a database of phylogenetic knowledge. In e-BioSphere 09: International Conference on Biodiversity Informatics, pages 216–217, London, UK. https://treebase.org/treebase-web/reference.html.

Posada, D. and Buckley, T. R. (2004). Model selection and model averaging in phylogenetics: advantages of Akaike Information Criterion and Bayesian approaches over likelihood ratio tests. Systematic Biology, 53:793–808.

Pupko, T. and Mayrose, I. (2020). A gentle introduction to probabilistic evolutionary models. In Scornavacca, C., Delsuc, F., and Galtier, N., editors, Phylogenetics in the Genomic Era, chapter 1.1, pages 1.1:1–1.1:21. Authors’ open access book. https://hal.archives-ouvertes.fr/hal-02535102.

Ronquist, F., Teslenko, M., van der Mark, P., Ayres, D. L., Darling, A., Höhna, S., Larget, B., Liu, L., Suchard, M. A., and Huelsenbeck, J. P. (2012). MrBayes 3.2: efficient Bayesian phylogenetic inference and model choice across a large model space. Systematic Biology, 61:539–542.

Soubrier, J., Steel, M., Lee, M. S., Der Sarkissian, C., Guindon, S., Ho, S. Y., and Cooper, A. (2012). The influence of rate heterogeneity among sites on the time dependence of molecular rates. Molecular Biology and Evolution, 29:3345–3358.

Uzzell, T. and Corbin, K. W. (1971). Fitting discrete probability distributions to evolutionary events. Science, 172:1089–1096.

Wong, T. K. F., Ly-Trong, N., Ren, H., Demotte, P., Baños, H., Roger, A. J., Susko, E., Bielow, C., De Maio, N., Goldman, N., Hahn, M. W., dos Reis, M., Vinh, L. S., Huttley, G., Lanfear, R., and Minh, B. Q. (2026). IQ-TREE 3: phylogenomic inference software using complex evolutionary models. Molecular Biology and Evolution, 43:msag117.

Yang, Z. (1993). Maximum-likelihood estimation of phylogeny from DNA sequences when substitution rates differ over sites. Molecular Biology and Evolution, 10:1396–1401.

Yang, Z. (1994). Maximum likelihood phylogenetic estimation from DNA sequences with variable rates over sites: approximate methods. Journal of Molecular Evolution, 39:306–314.

Yang, Z. (1995). A space-time process model for the evolution of DNA sequences. Genetics, 139:993–1005.

Yang, Z. (1996). Among-site rate variation and its impact on phylogenetic analyses. Trends in Ecology & Evolution, 11:367–372.

Yang, Z. (2007). PAML 4: Phylogenetic Analysis by Maximum Likelihood. Molecular Biology and Evolution, 24:1586–1591.

Yang, Z. (2010). A likelihood ratio test of speciation with gene flow using genomic sequence data. Genome Biology and Evolution, 2:200–211.

Yang, Z., Nielsen, R., Goldman, N., and Pedersen, A.-M. K. (2000). Codon-substitution models for heterogeneous selection pressure at amino acid sites. Genetics, 155:431–449.

Yang, Z. and Roberts, D. (1995). On the use of nucleic acid sequences to infer early branchings in the tree of life. Molecular Biology and Evolution, 12:451–458.

